# Disease-Associated Non-Coding Variants Alter NKX2-5 DNA-Binding Affinity

**DOI:** 10.1101/2022.12.02.518772

**Authors:** Edwin G. Peña-Martínez, Alejandro Rivera-Madera, Diego A. Pomales-Matos, Leandro Sanabria-Alberto, Brittany M. Rosario-Cañuelas, Jessica M. Rodríguez-Ríos, Emmanuel A. Carrasquillo-Dones, José A. Rodríguez-Martínez

**Affiliations:** University of Puerto Rico-Río Piedras Campus, San Juan, Puerto Rico; University of Puerto Rico-Cayey Campus, Cayey, Puerto Rico

**Keywords:** transcription factors, non-coding variants, binding affinity, gene regulation

## Abstract

1.

Genome-wide association studies (GWAS) have mapped over 90% of disease- or trait-associated variants within the non-coding genome, like *cis*-regulatory elements (CREs). Non-coding single nucleotide polymorphisms (SNPs) are genomic variants that can change how DNA-binding regulatory proteins, like transcription factors (TFs), interact with the genome and regulate gene expression. NKX2-5 is a TF essential for proper heart development, and mutations affecting its function have been associated with congenital heart diseases (CHDs). However, establishing a causal mechanism between non-coding genomic variants and human disease remains challenging. To address this challenge, we identified 8,475 SNPs predicted to alter NKX2-5 DNA-binding using a position weight matrix (PWM)-based predictive model. Five variants were prioritized for in vitro validation; four of them are associated with traits and diseases that impact cardiovascular health. The impact of these variants on NKX2-5 binding was evaluated with electrophoretic mobility shift assay (EMSA) using recombinantly expressed and purified human NKX2-5 homeodomain. Binding curves were constructed to determine changes in binding between variant and reference alleles. Variants rs7350789, rs7719885, rs747334, and rs3892630 increased binding affinity, whereas rs61216514 decreased binding by NKX2-5 when compared to the reference genome. Our findings suggest that differential TF-DNA binding affinity can be key in establishing a causal mechanism of pathogenic variants.

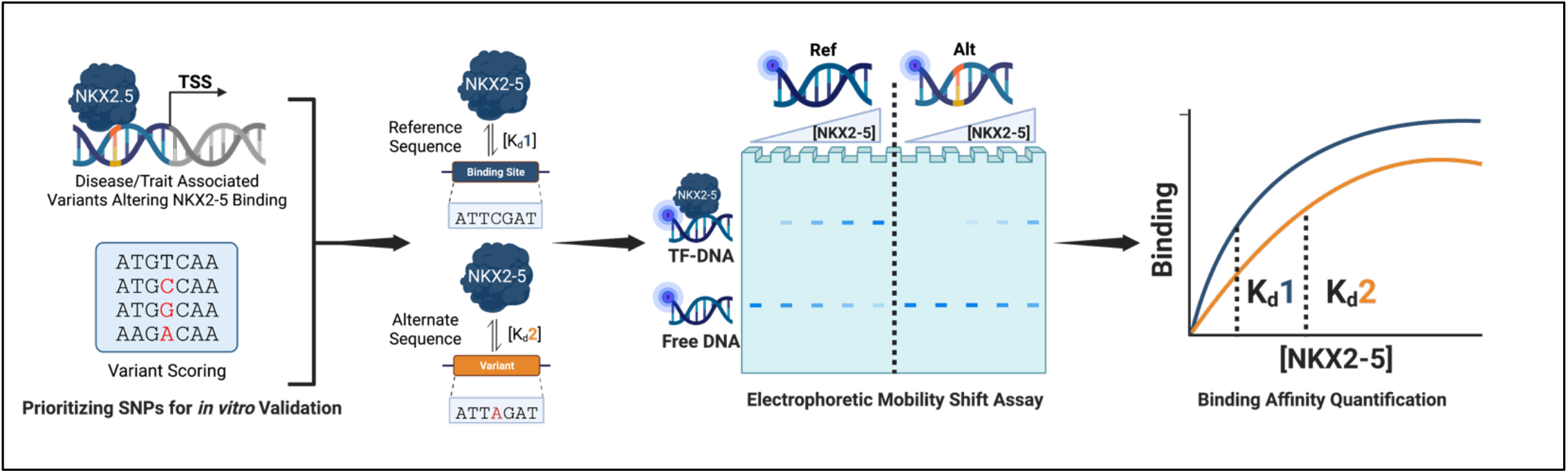

## 2. Introduction

Genome-wide association studies (GWAS) have revealed that over 90% of disease/trait-associated variants occur within the non-coding genome. ^1–5^ Non-coding DNA comprises 98% of the human genome and includes regions like *cis*-regulatory elements (CREs), such as promoters and enhancers, that are essential for regulating gene expression.^6–11^ Non-coding single nucleotide polymorphisms (SNPs) within CREs can alter the function of regulatory DNA-binding proteins, like transcription factors (TFs). ^12–14^ TFs bind to specific DNA sequences within CREs and recruit chromatin remodelers and the transcriptional machinery to regulate gene expression. ^15–19^ Non-coding SNPs can result in binding affinity changes between a TF and its binding site, leading to dysregulation of biological processes like cell growth, differentiation, and organ development. ^20–24^

NKX2-5 is an evolutionarily conserved tissue-specific TF involved in heart development. This cardiac TF is essential in the gene regulatory network for myocyte differentiation and cardiac morphogenesis. ^25–32^ Previous work has proven mutations within the NKX2-5 DNA-binding domain (DBD), a homeodomain, can alter its regulatory function, leading to cardiovascular diseases like congenital heart defects (CHDs). ^33–35^ However, most variants associated with cardiovascular disease are non-coding and occur in CREs that can function as cardiac TF binding sites (TFBS). ^36–39^ Non-coding SNPs can affect TF-DNA binding affinity resulting in the loss of TFBS or the creation of a new TFBS in the genome ^39–41^. Previous work has shown that coronary artery disease (CAD)-associated SNPs have disrupted the binding of tissue-specific TFs like STAT1, MEF2, and KLF2.^42–44^

In this work, we evaluated the potential of disease-associated variants to alter the binding affinity of the cardiac TF NKX2-5. Using SNP2TFBS^45^, a position weight matrix (PWM)-based predictive model, we identified five variants associated with cardiovascular-related traits predicted to impact NKX2-5 binding. Through electrophoretic mobility shift assays (EMSA), we observed differential NKX2-5 binding activity between the reference and alternate genomic sequences. Binding curves were constructed to derive apparent dissociation constants (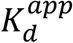) and determine changes in NKX2-5 binding. We found that three out of five predictions were consistent with *in vitro* observations, whereas for the other two variants *in vitro* experiments contradicted the *in silico* prediction. Our results show that non-coding cardiovascular trait-associated SNPs change NKX2-5 DNA binding affinity. This suggests that disruption or creation of TFBS throughout the genome can be a plausible causal mechanism behind cardiovascular diseases like CADs and CHDs.

## 3. Materials and Methods

### 3.1 Identification of Non-coding Variants

Non-coding single nucleotide polymorphisms (SNPs) predicted to alter NKX2-5-DNA binding affinity were identified using SNP2TFBS, a position weight matrix (PWM)-based predictive model. ^45^ Using SNP2TFBS, we used an NKX2-5 PWM (MA0063.2) from the JASPAR Core 2014 vertebrates^46^ database to score variants cataloged from the 1000 Genomes Project^47^. Variants predicted to alter NKX2-5 binding were intersected with SNPs in the GWAS catalog^3^. Filtered variants were scored using the SNP2TFBS, and the top five variants with the largest predicted change in binding were selected for *in vitro* validation.

### 3.2 NKX2-5 cloning and expression

The NKX2-5 homeodomain gene (DNASU Plasmid Repository) was cloned in a pET-51(+) expression vector (Novagen) containing an N-terminal Strep•Tag^®^ and a C-terminal 10X His•Tag^®^ through Gibson Cloning and used to transform BL21 DE3 *E. coli* strain (Millipore). Glycerol stock of transformed bacteria was cultivated in 50 mL Luria Broth (Sigma-Aldrich) for 16 h at 37°C. Following this period, 10 mL of initial bacterial culture was transferred to 500 mL of Terrific Broth (Sigma-Aldrich) in a 2 L Erlenmeyer flask and grown at 37°C at 130 rpm until the optical density at 600 nm (OD600) reached 0.5. Once the OD600 reached 0.5, the bacterial culture was induced with 1 mM IPTG for 20 h at 20°C and shaking at 130 rpm. The cells were collected by centrifugation (2,800 x g, 5 min, 4°C), the supernatant was discarded, and the pellet was stored at -80°C overnight.

### 3.3 Protein purification

NKX2-5 HD was purified from lysed bacterial cells through Ni-NTA affinity chromatography. Cell pellets were resuspended in 40 mL column buffer (500 mM NaCl, 20 mM Tris-HCl (pH 8.0), 0.2% Tween-20, 30 mM imidazole, and EDTA-free protease inhibitor). 4 mL 5 M NaCl were added to resuspended cells and sonicated in four 30-second cycles at 40% amplitude (QSONICA, Part No. Q125). The lysate was centrifuged (2,800 x g, 30 min, 4°C), and the supernatant was loaded into the Ni-NTA affinity chromatography column. The column was prepared with 2 mL Ni-NTA Agarose Resin (Qiagen). The resin was equilibrated with 10 column volumes of column buffer and resuspended with the supernatant cell extract for 1 h at 4°C with orbital shaking. Column flowthrough was passed again through the column twice. The column was washed with 20 mL of column buffer thrice, with increasing the imidazole concentration (30, 50, and 100 mM) in each wash. The column was eluted with 1.8 mL of elution buffer (500 mM NaCl, 20 mM Tris-HCl (pH 8.0), 0.2% Tween-20, 500 mM imidazole) six times.

NKX2-5 homeodomain purity was evaluated through SDS-PAGE using 15 well Mini-PROTEAN TGX Precast Protein Gels^®^ (Bio-Rad). Samples were prepared with 4X loading buffer containing β-mercaptoethanol (BME) for a total volume of 20 μL (5μL 4x loading buffer: 15 μL sample) and heated at 95°C for 5 min. 15 μL were loaded onto the gel for a 1.5 h run at 100 V at room temperature. The gel was stained using ProtoStain™ Blue Colloidal Coomasie G-250 stain (National Diagnostics). For the Western Blot analysis, contents from the SDS-PAGE gel were transferred to a PVDF membrane using the Bio-Rad Turbo Transfer System protocol in a Trans-Blot^®^ Turbo™ for 3 min at 25 V. Membranes were blocked using 5% milk in TBST buffer for 1 h in orbital shaking and incubated overnight with 1:10,000 dilution of Anti-His mouse monoclonal antibodies (Novus Biologicals, AD1.1.10). The SDS-PAGE and Western Blot Analysis results were imaged using Azure^®^ Biosystems Imager.

### 3.4 Electrophoretic Mobility Shift Assay

NKX2-5 binding was evaluated using 40 bp sequences centered on the SNP and an additional 20 bp sequence for IR-700 fluorescent marking (IDT). All sequences were ordered in IDT and are available in **Supplementary Table 2**. Reference and alternate sequences were added the IR-700 fluorophore through a primer extension reaction. Binding reactions were performed in binding buffer (50 mM NaCl, 10 mM Tris-HCl (pH 8.0), and 10% glycerol) and 5 nM fluorescently labeled dsDNA. Binding reactions were incubated for 30 min at 30°C and 30 min at room temperature before loading onto a 6% polyacrylamide gel in 0.5x TBE (89 mM Tris/89 mM boric acid/2 mM EDTA, pH 8.4). The gel was pre-runned at 85 V for 15 min, loaded at 30 V, and resolved at 75 V for 1.5 h at 4°C. Gels were imaged with Azure^®^ Sapphire Bio-molecular Imager at 658 nm/710 nm excitation and emission.

### 3.5 Binding Affinity

Apparent K_d_ was determined by first quantifying the fluorescence signal in each DNA band using ImageJ^48^. Background intensities obtained from blank regions of the gel were subtracted from the band intensities. The fraction of bound DNA was determined using **Equation 1**. The fraction of bound DNA was plotted versus the concentration of the NKX2-5 homeodomain. Binding curves, K_d_, and B_max_ were obtained by “one-site specific binding” non-linear regression using Prism software.

**Equation 1**. Binding Affinity from the integrated density of bound and unbound bands.

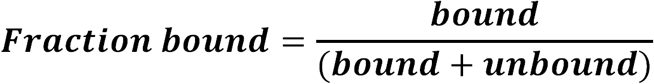

## 4. Results and Discussion

### 4.1 Identification of Non-Coding Disease-Associated SNPs

Using a PWM-based predictive model, we identified 8,475 SNPs predicted to change NKX2-5 binding out of the >84 million SNPs cataloged from the 1000 Genomes project. ^47^ The genomic coordinates of the predicted variants were intersected with disease or quantitative trait-associated SNPs from the GWAS catalog, resulting in 30 variants (**Figure 1**). The output of the SNP2TFBS includes a ΔPWM score that predicts if the TF-DNA binding will increase or decrease based on a positive or negative score, respectively. ΔPWM scores were sorted by magnitude, and the five variants with the largest predicted impact on NKX2-5 DNA binding were chosen for *in vitro* validation, including two variants with a predicted increase in binding and three variants with decreased binding (**Table 1**). The selected SNPs are associated with traits and diseases that impact cardiovascular phenotypes (e.g., hemoglobin and cholesterol levels, red blood cell traits, and systolic blood pressure).

**Table 1:** Non-coding SNPs prioritized for in vitro validation.

| rsID | Mutation | ΔPWM Score | Predicted Binding Impact | Associated Disease or Trait |
| --- | --- | --- | --- | --- |
| rs7350789 | G → A | 258 | Increase | Serum metabolite levels, High density lipoprotein cholesterol levels, Postprandial triglyceride response, Total cholesterol levels, Phosphatidylethanolamine levels |
| rs61216514 | G → A | -232 | Decrease | Mean corpuscular hemoglobin |
| rs7719885 | A → G | -212 | Decrease | Systolic blood pressure |
| rs747334 | A → G | -187 | Decrease | Fibroblast growth factor basic levels |
| rs3892630 | C → G | 146 | Increase | Red blood cell traits |

**Figure 1:**
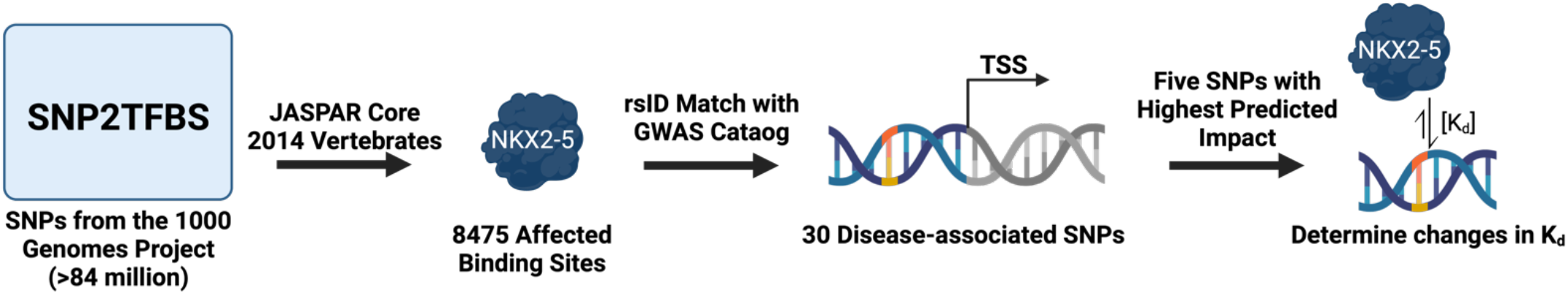
Identification of disease/trait-associated non-coding SNPs affecting NKX2-5 binding. Predicted SNPs from the 1000 Genomes Project were intersected with disease-associated variants from the GWAS catalog.

### 4.2 Expression and Purification of NKX2-5 Homeodomain

Recombinant NKX2-5 homeodomain with an N-terminal Strep•Tag^®^ and a C-terminal 10X His•Tag^®^ was cloned in an expression vector and produced using an IPTG-inducible bacterial system. After overexpression of NKX2-5 homeodomain, bacteria were lysed and centrifuged to obtain soluble fractions and purified using Ni-NTA affinity chromatography and eluted with an imidazole gradient. The NKX2-5 homeodomain was successfully purified as determined by SDS-PAGE and Western Blot (**Figure 2A; Supplementary Figure 1**). The DNA-binding activity of the purified NKX2-5 homeodomain was determined through EMSA using a known binding site within the ANF promoter (**Figure 2B-C; Supplementary Figure 2**).

**Figure 2:**
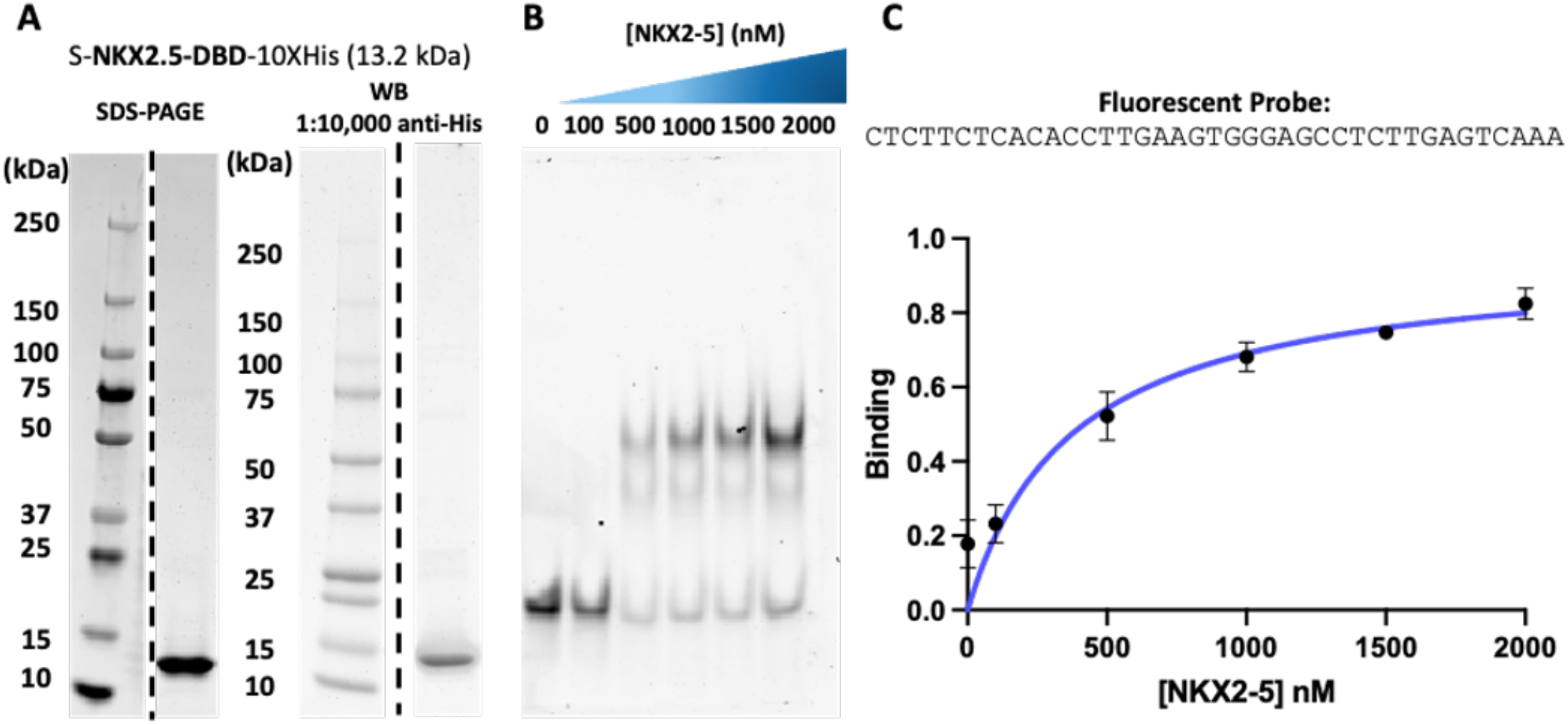
Expression and purification of functional NKX2-5 homeodomain. A) SDS-PAGE (left) and Western blot (right) of purified NKX2-5 homeodomain after Ni-NTA affinity chromatography. B) Electrophoretic mobility shift assay (EMSA) of known binding site within the ANF promoter. C) Binding curve analysis of NKX2-5 homeodomain.

### 4.3 Non-coding mutations alter NKX2-5 binding

We tested the five variants with the largest predicted change in binding using 40 bp of genomic sequences centered at the SNP. Oligonucleotides were synthesized with an additional 20 bp constant region that served to add the fluorescent probe via primer extension. Changes in DNA binding affinity between the reference and the alternate allele were determined through EMSA. Purified NKX2-5 homeodomain was equilibrated with reference and variant sequences at seven different protein concentrations. Differential TF-DNA binding was observed for the five predicted variants (**Supplementary Figure 3**). The impact of the five variants on binding affinity was quantified by generating binding curves and calculating the 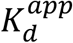values (**Figure 3**). Three out of the five tested variants agreed with our computational prediction (**Supplementary Table 1**). Variants rs7350789 and rs3892630 were precited by SNP2TFBS to increase binding affinity and were successfully validated by EMSA. Changes in 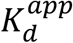 for rs3892630 resulted in a 2.3-fold decrease, while rs7350789 could not be quantified due to low binding affinity in the reference sequence. SNP2TFBS predicted variant rs61216514 to decrease in binding affinity and was successfully validated by EMSA, resulting in a 1.3-fold change increase of its 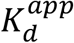. However, the two variants that did not follow the prediction (rs7719885 and rs747334) still demonstrated differential DNA binding. SNP2TFBS predicted variants rs7719885 and rs747334 would decrease binding affinity. Both increased, resulting in a 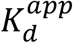 1.3- and 1.8-fold change decrease, respectively. The predictions are made using a PWM model generated from chromatin immunoprecipitation sequencing (ChIP-seq) data which has cellular factors not present in a biochemical assay like an EMSA. Additional DNA-binding specificity models generated by *in vitro* experiments or alternate computational approaches will be tested in the future. However, all five variants significantly altered NKX2-5 DNA binding affinities as determined by changes in their 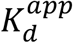 values.

**Figure 3:**
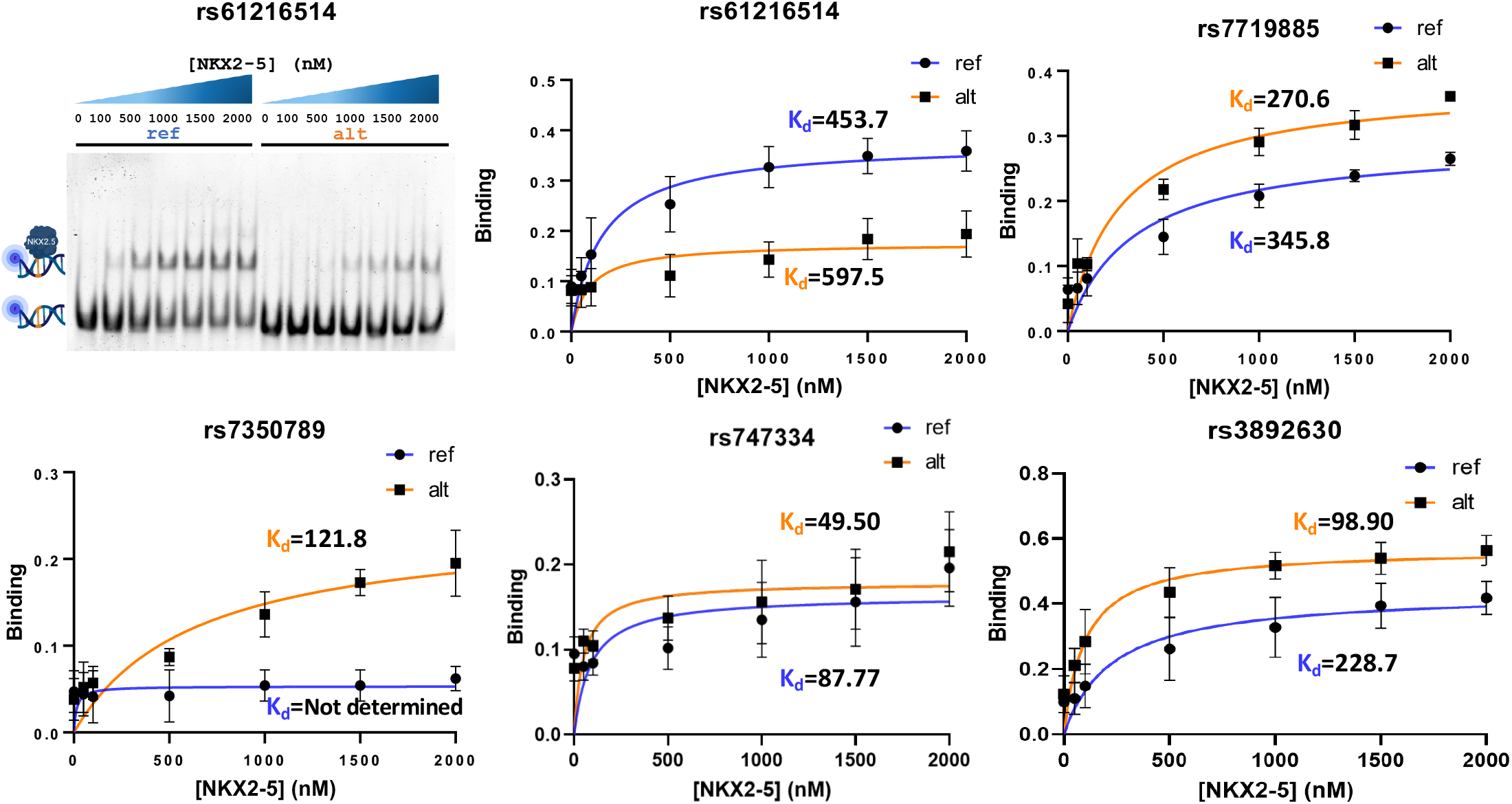
NKX2-5 binding to reference (ref) and variant (alt) sequences determined through EMSA (triplicates). Gel electrophoresis image used for binding curve analysis for variants rs6121514 is shown (top left).

## 5. Conclusion

We conclude that non-coding GWAS variants can alter NKX2-5 affinity for its genomic binding sites. Non-coding SNPs associated with cardiovascular traits altered the NKX2-5-DNA complex formation, which is essential for heart development. Changes in the biophysical properties of gene regulation, like TF-DNA binding, are key factors to consider when determining the causal mechanism of genetic variants behind human diseases. Pathogenic SNPs within the NKX2-5 binding sites are potential regulatory targets for healthy cardiovascular development and function.

## 6. Acknowledgments

We thank Dr. Esther Peterson for providing the software license to graph our results. This project was supported by NIH-SC1GM127231. EGPM and JMRR were funded by the NSF BioXFEL Fellowship (STC-1231306). EGPM and DAPM were funded by the NIH RISE Fellowship (5R25GM061151-20). ARM was funded by NSF REU: PR-CLIMB Program (2050493). LSA was funded by NIH IG-GENE Fellowship (1R25HG012702-01). JMRR was funded NSF Graduate Research Fellowship (1744619). BMRC was funded by the ACS SEED Program and the UPRRP Department of Chemistry. Graphical abstract and Figure 1 were created in Biorender^®^.

## 8. Supplementary material

**Supplementary Figure 1:**
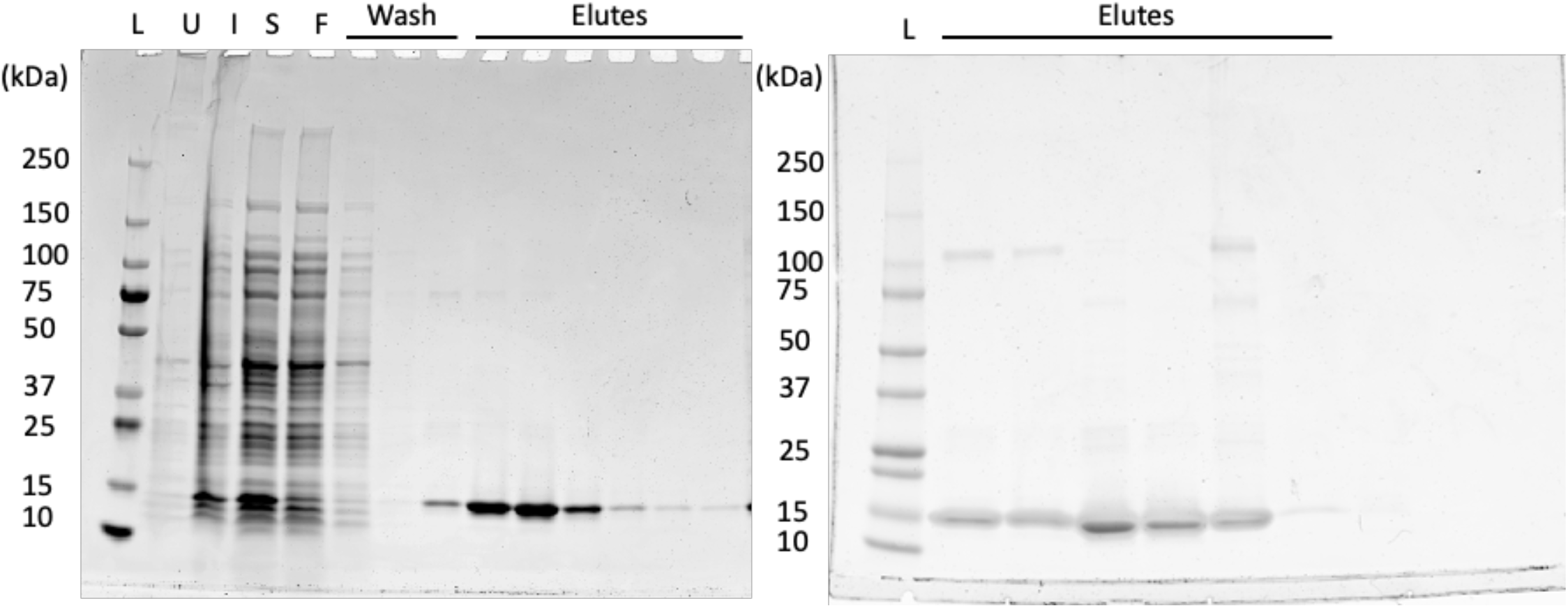
(left) SDS-PAGE analysis of Ni-NTA affinity chromatography. NKX2-5 homeodomain purification fractions: ladder (L), uninduced (U), induced (I), supernatant (S), flowthrough (F), imidazole gradient wash (30, 50, and 100 nM) and protein elutes. (right) Western Blot identification of NKX2-5 homeodomain elutes.

**Supplementary Figure 2:**
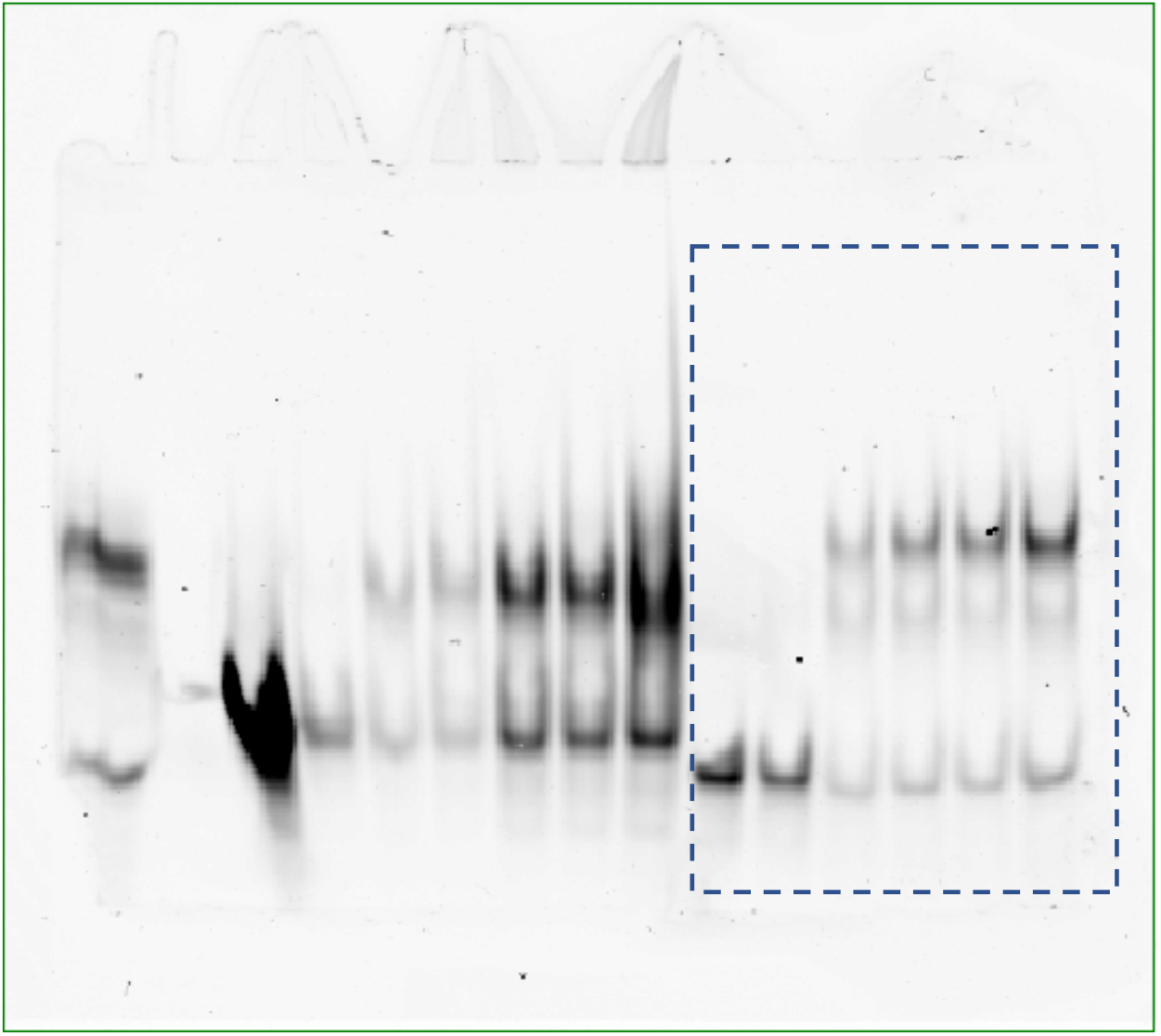
EMSA analysis of functional NKX2-5 homeodomain at 0, 100, 500, 1000, 1500, and 2,000 nM. Bands used in Figure 2B are inside the dashed square.

**Supplementary Figure 3:**
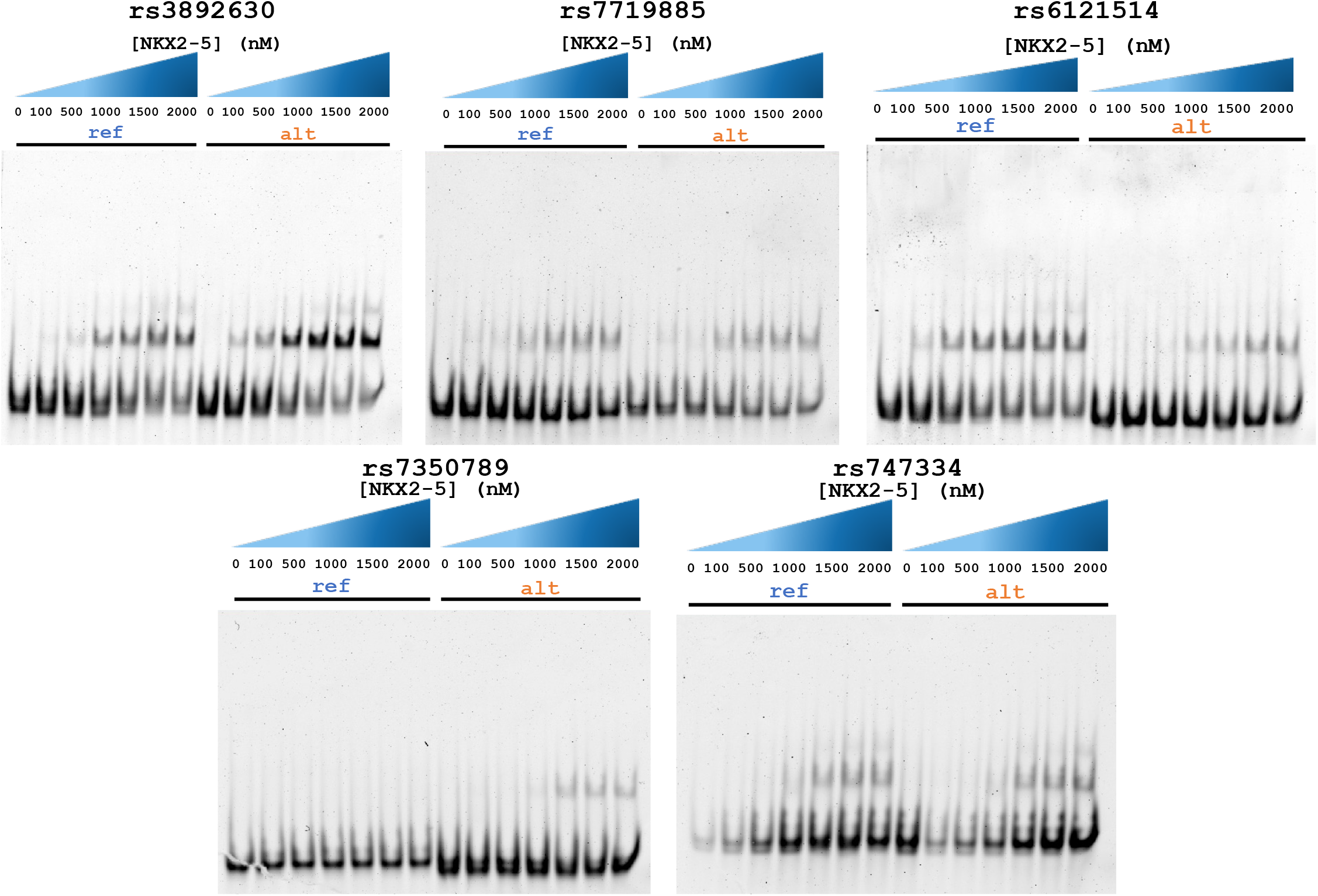
EMSA analysis of five disease-associated non-coding SNPs.

**Supplementary Table 1:**
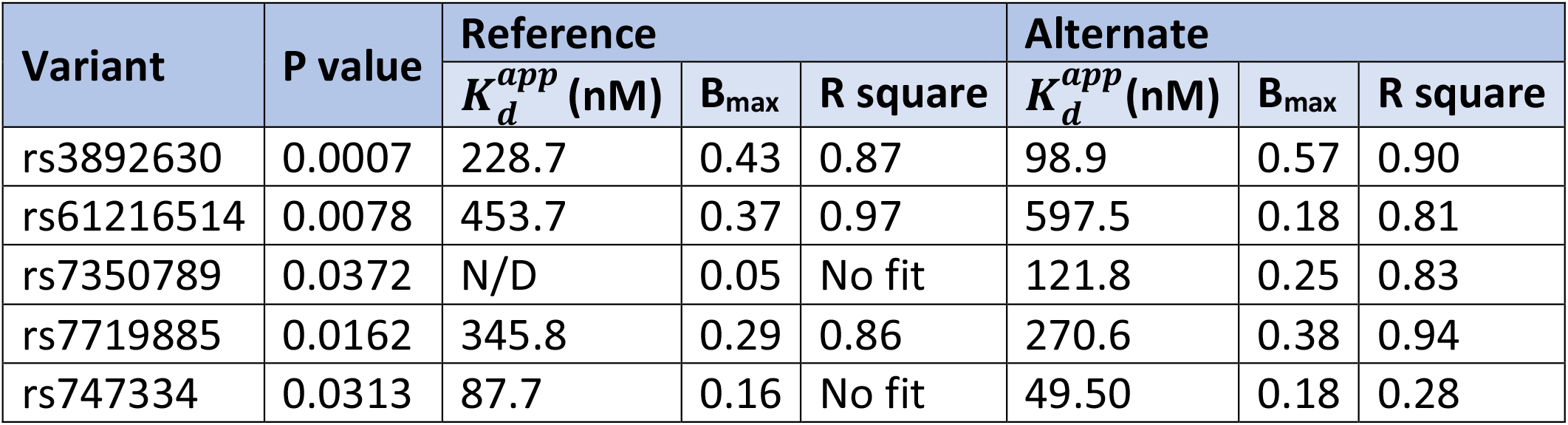
Non-lineal regression parameters of the five disease-associated non-coding SNPs.

**Supplementary Table 2:**
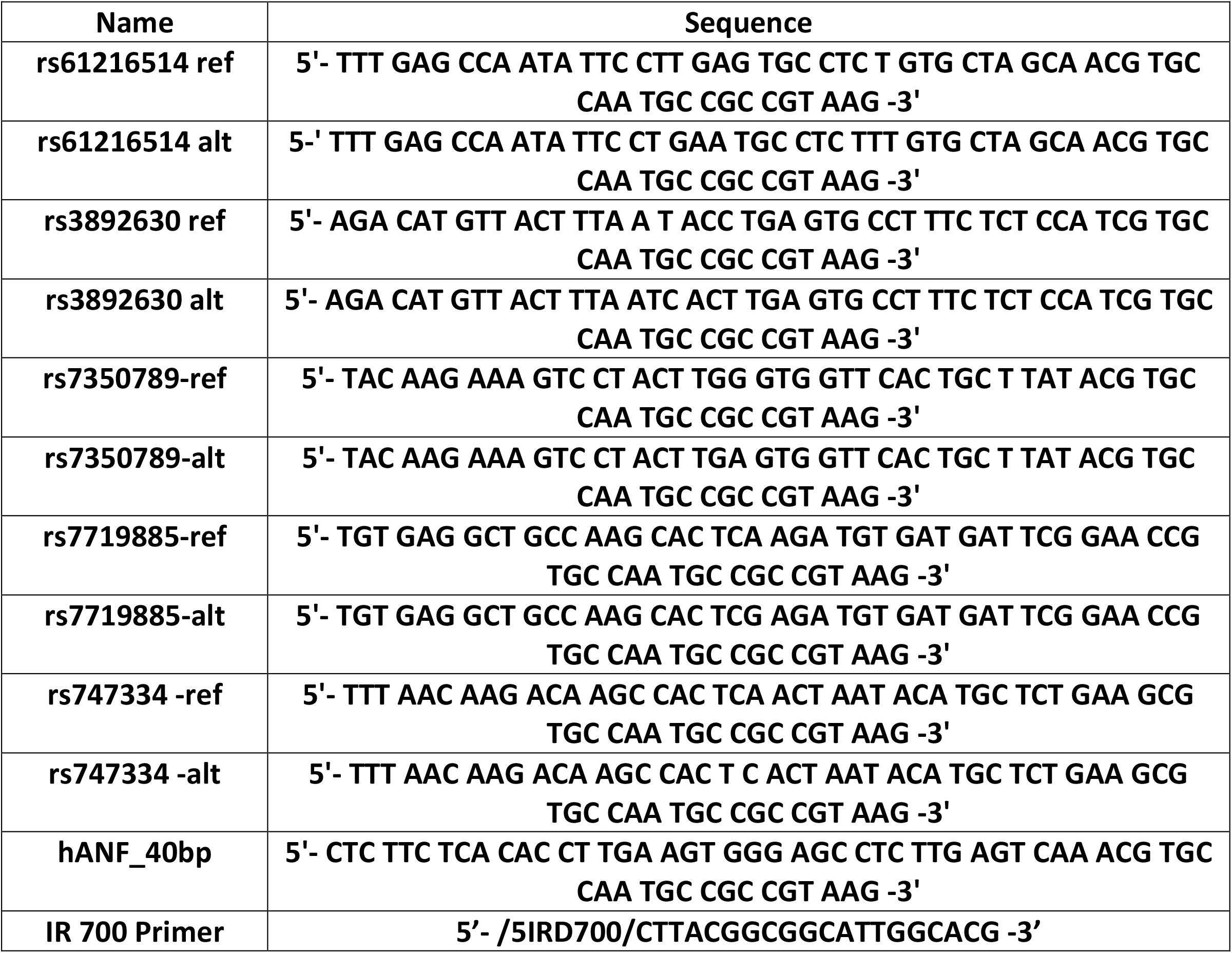
List of sequences.

